# xQTLImp: efficient and accurate xQTL summary statistics imputation

**DOI:** 10.1101/726182

**Authors:** Tao Wang, Quanwei Yin, Yongzhuang Liu, Jin Chen, Yadong Wang, Jiajie Peng

## Abstract

**Motivation:** Quantitative trait locus (QTL) analysis of multiomic molecular traits, such as gene transcription (eQTL), DNA methylation (mQTL) and histone modification (haQTL), has been widely used to infer the effects of genomic variation on multiple levels of molecular activities. However, the power of xQTL (various types of QTLs) detection is largely limited by missing association statistics due to missing genotypes and limited effective sample size. Existing hidden Markov model (HMM)-based imputation approaches require individual-level genotypes and molecular traits, which are rarely available. No available implementation exists for the imputation of xQTL summary statistics when individual-level data are missed.

**Results:** We present xQTLImp, a C++ software package specifically designed for efficient imputation of xQTL summary statistics based on multivariate Gaussian approximation. Experiments on a single-cell eQTL dataset demonstrates that a considerable amount of novel significant eQTL associations can be rediscovered by xQTLImp.

**Availability:** Software is available at https://github.com/hitbc/xQTLimp.

**Supplementary information:** Supplementary data are available at *Bioinformatics* online.

## 1 Introduction

Quantitative trait locus (QTL) analyses of multiomic phenotypes are critical in understanding the complex functional regulatory natures of disease-associated variants unequivocally identified by genome wide association studies (GWAS) (Ng *et al.*, 2017). The summary statistics of xQTL studies, such as eQTL, mQTL and haQTL, have been widely used in estimating causal effects of GWAS variants, such as in mendelian randomization analysis or meta-analysis, to boost the statistical power (GTEx Consortium, 2017). However, one critical problem here is the missing of xQTL statistics in every individual study, which may bias the findings and generate many false negative results because of power lost (Furukawa *et al.*, 2006). The missing statistics can be attributed to two main reasons. First, missing genotypes are common in various studies because of different genotyping platforms, genotype imputation methods, or quality control procedures. Although excellent hidden Markov model (HMM)-based genotype imputation tools exist, these tools require individual-level genotypes, which are typically hard to access in practice due to ethical restrictions (Marchini and Howie, 2010). Second, xQTL analysis requires both genotypes and molecular traits for the same individual, which might cause a limited effective sample size due to factors such as cost or sample availability. A small sample size will limit the minor allele frequency (MAF) lower bound in association tests, which will cause missing statistics even if the genotype is present. For example, the recent single-cell eQTL studies can only perform eQTL analyses on variants with a MAF ≥ 0.1 (van der Wijst *et al.*, 2018) and 0.2 (Kang *et al.*, 2018) with effective sample sizes of 45 and 23, respectively.

To the best of our knowledge, there is no existing tool that is designed for the imputation of xQTL summary statistics. To fill this gap, we developed xQTLImp, a C++ tool designed specifically for efficiently imputing xQTL summary statistics. xQTLImp models the statistics of variants in linkage disequilibrium (LD) associated with the same molecular trait by using a multivariate Gaussian model (Supplementary materials), which has been successfully applied in the GWAS era, such as multiple correlated marker correction (Han *et al.*, 2009), fine mapping (Zaitlen *et al.*, 2010) and GWAS imputation (Wen and Stephens, 2010; Pasaniuc *et al.*, 2014; Kwan *et al.*, 2016). Experiments on real eQTL, mQTL and haQTL datasets demonstrated the high imputation accuracy of xQTLImp and ability to discover novel signals which will enhance discoveries in xQTL studies.

## 2 Methods

The details of the methods can be found in the supplementary notes. In brief, given a molecular trait *G* and cis-variants around *G* in a reference panel, the objective of xQTLImp is to impute unknown variant-*G* association statistics, from incomplete user-given variant-*G* association statistics by leveraging LD among markers. xQTLImp requires as compulsory input data: (1) genotype reference panel for calculating LD correlations; (2) xQTL summary statistics (Z-scores of variant-G associations), where variants are identified by genomic coordinates and ref/alt alleles; (3) molecular trait annotation. xQTLImp also allows users to specify the chromosome and MAF of variants, discard certain regions and run in multiple threads. We model xQTL summary statistics (Z-scores) by using multivariate normal (MVN) distribution. Formally, Let ***Z***_***u***_|*G* and ***Z***_***k***_|*G* be two vectors with lengths *n* and *m*, representing unknown and known xQTL statistics of *n* and *m* variants associated with trait G, respectively. Our objective is to impute ***Z***_***u***_|*G* from ***Z***_***k***_|*G* given known LD matrix Σ, representing the LD correlations among *n* + *m* variants. We estimate 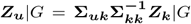, derived by conditional MVN distribution of ***Z***_***u***_|*G* given ***Z***_***k***_|*G*. We adopt an imputation quality score *r*2*pred* ∈ [0, 1) to measure the xQTL imputation accuracy.

## 3 Results

### 3.1 Imputation accuracy

To test the imputation accuracy of xQTLImp, we first randomly masked certain percentage of summary statistics of eQTL from GTEx study v7 (brain Amygdala, *n* = 88) (GTEx Consortium, 2017), haQTL and mQTL from the brain xQTL study (frontal cortex, *n* = 433 and 468, respectively) (Ng *et al.*, 2017) on chromosome 1, with MAF ≥ 0.01 for all variants. Second, xQTLImp was applied to impute the concealed statistics based on 1000G phase3 reference panel (EUR population, *n* = 503) (1000 Genomes Project Consortium, 2015). The results in Supplementary Fig. 1 show that xQTLImp achieves high imputation accuracy with a Pearson’s *r >* 0.95 in all masking percentages under the thresholds of *r*2*pred* ≥ 0.6. Specifically, more than 90% masked associations can be recovered by imputation with *r*2*pred* ≥ 0.6 (Supplementary Fig. 1). Similar results were observed in other chromosomes (Supplementary Fig. 2).

Since xQTLImp does not require individual-level datasets, it has the potential to impute novel associations with MAF lower than the lower bound in the original xQTL studies. To test the imputation accuracy on lower MAF levels, we adopted two WGS-based eQTL datasets from GTEx study v7 (brain amygdala and muscle skeletal, *n* = 88 and 491, respectively). eQTL associations with MAF *>* 0.1 and MAF *>* 0.05 were used to impute associations with 0.05 ≤ MAF ≤ 0.1 and 0.01 ≤ MAF ≤ 0.05 in brain amygdala and muscle skeletal, respectively, using the same reference panel as above. As a result, xQTLImp achieves a high accuracy with Pearson’s r = 0.95, 0.92 respectively by setting a stringent r2pred cutoff as 0.9 (Supplementary Fig. 3).

For performance comparison, we also compared xQTLImp with HMM-based genotype imputation framework, which is always regarded as the gold standard but require individual-level genotype which are typically hard to access in practice. We performed eQTL analysis using individual-level genotypes and RNA-seq profiles from the ROSMAP study (Mostafavi *et al.*, 2018) after stringent quality controls (Supplementary Notes), and compared the imputation performance between the two strategies. The result showed a high consistency (Pearson’s r = 0.996) between statistics from xQTLImp imputation and HMM-based imputation (Supplementary Fig. 4), which indicates that xQTLImp can accurately obtain xQTL statistics without individual-level data.

### 3.2 Case study on single-cell eQTL datasets

We applied xQTLImp to summary statistics from a single-cell RNA-sequencing-based eQTL study with six major cell types (CD4+ T cells, CD8+ T cells, natural killer (NK) cells, monocytes, dendritic cells (DCs) and B cells) detected from peripheral blood mononuclear cells (PBMCs) from 45 donors (van der Wijst *et al.*, 2018). The MAF lower bound was 0.1 in the original study due to a limited sample size. We used xQTLImp to impute novel eQTL associations with MAF ≥ 0.01 using a 1000G EUR reference panel. By applying a stringent r2pred cutoff of 0.9 and the same permutation-based significance P-value cutoffs (FDR < 0.05) as in the original study, we successfully detected a large amount of novel significant eQTL associations (an increase of 64% to 148% compared with original significant findings) (Supplementary Table 1). A considerable number of novel significant cell-type-specific eQTL associations were also revealed (Supplementary Fig. 5). Both original significant findings and novel significant findings can be well replicated in an independent CD4+ T cell-type-specific eQTL dataset (Supplementary Fig. 6).

Regarding the performance, xQTLImp took 3 ∼ 4 hours and <4 GB memory for whole genome-wide imputation on each cell-type in the discovery study. It used 20 threads and the other settings as their default values (Supplementary Table 2).

## 4 Conclusion

In conclusion, we present xQTLImp, an efficient and accurate tool designed specifically for xQTL summary statistics imputation. Using multiple real datasets, we demonstrated that 1) xQTLImp can impute missing xQTL statistics with high accuracy, and 2) xQTLImp can effiectively reduce the lower bound of MAF in xQTL studies, leading to the discovery of novel xQTL signals to further enhance xQTL discoveries.

