## Supplementary material for "xQTLImp: efficient and accurate xQTL summary statistics imputation": online

#### Supplementary Materials

Tao Wang[1,†], Quanwei Yin[2,3,†], Yongzhuang Liu[1], Jin Chen[4], Yadong Wang[1,*], and Jiajie Peng[2,3,*]

[1]School of Computer Science and Technology, Harbin Institute of Technology, Harbin, 150001, China
[2]School of Computer Science, Northwestern Polytechnical University, Xi'an, 710072, China
[3]Key Laboratory of Big Data Storage and Management, Ministry of Industry and Information Technology, Northwestern Polytechnical University, Xi'an, 710072, China
[4]Institute for Biomedical Informatics, University of Kentucky, Lexington, KY 40536, USA

4 Dataset availability 5
5 Acknowledgment 5
6 Supplementary Figures 6
7 Supplementary Tables 12

Figure S6 Diagram of two-way contingency table for Fisher's exact test. 11

† *Equal contributor.*

### LIST OF TABLES

Table S1 Summary of novel significant eQTL associations found by xQTLImp compared with that in original single-cell eQTL datasets of six cell types. . . . . . . . . . . . . . . . . . . . . . . . 12

Table S2 Running time and memory cost on single-cell eQTL datasets of six cell types. . . . . . . . . . . . . . . . . . . . . . . . . . . . 13

#### 1 STATISTICAL MODEL

Similar to the statistical model described in GWAS imputation works [1, 2, 3, 4], we model the xQTL summary statistics, given by Z-scores, of variants in association with molecular trait G by using a multivariate normal distribution (MVN). Suppose that $n+m$ variants are associated with molecular trait G and that Z scores are represented as $\mathbf{Z}|G = (x_1, ..., x_n, y_1, ..., y_m)^T$, where $\mathbf{Z_u}|G = (x_1, ..., x_n)^T$ represents unknown xQTL statistics of $n$ variants associated with G, which needs to be imputed, and $\mathbf{Z_k}|G = (y_1, ..., y_m)^T$ represents known xQTL statistics of $m$ variants associated with G. We model $\mathbf{Z}|G$ by MVN, i.e., $\mathbf{Z}|G \sim N(\boldsymbol{\mu}, \boldsymbol{\Sigma})$, where $\boldsymbol{\mu}$ represents the vector of means with length $n+m$ and $\boldsymbol{\Sigma}$ represents the covariance matrix of $n+m$ variants with $\boldsymbol{\Sigma}_{i,j}$ representing the LD correlation $r$ between $SNP_i$ and $SNP_j$, which can be calculated from reference panels such as 1000G and HapMap. By denoting expectations $E(\mathbf{Z_u}|G) = \boldsymbol{\mu_u}$ and $E(\mathbf{Z_k}|G) = \boldsymbol{\mu_k}$, we have $\boldsymbol{\mu} = (\boldsymbol{\mu_u}, \boldsymbol{\mu_k})^T$, and similarly, $\boldsymbol{\Sigma}$ can be partitioned into four submatrices based on two groups of variants, i.e., $\boldsymbol{\Sigma} = \begin{bmatrix} \boldsymbol{\Sigma_{uu}} & \boldsymbol{\Sigma_{uk}} \\ \boldsymbol{\Sigma_{ku}} & \boldsymbol{\Sigma_{kk}} \end{bmatrix}$, where $\boldsymbol{\Sigma_{ku}} = \boldsymbol{\Sigma_{uk}^T}$.

Our objective is to impute $\mathbf{Z_u}|G$ from $\mathbf{Z_k}|G$ given a known LD matrix $\boldsymbol{\Sigma}$. This problem can be solved by the conditional distribution of $\mathbf{Z_u}|G$ given $\mathbf{Z_k}|G$, represented as $(\mathbf{Z_u}|\mathbf{Z_k})|G$. It has been proven that the conditional distribution for $\mathbf{Z_u}|G$ given $\mathbf{Z_k}|G$ is $n$-dimensional normal with expectation and covariance matrix as follows:

$$\begin{aligned} E[(\mathbf{Z_u}|\mathbf{Z_k})|G] &= \boldsymbol{\mu}_{Z_u|Z_k} = \boldsymbol{\mu_u} + \boldsymbol{\Sigma_{uk}}\boldsymbol{\Sigma_{kk}^{-1}}(Z_k|G - \boldsymbol{\mu_k}) \\ &= \boldsymbol{\Sigma_{uk}}\boldsymbol{\Sigma_{kk}^{-1}} Z_k|G \end{aligned} \tag{1}$$

$$\begin{aligned} Cov[(\mathbf{Z_u}|\mathbf{Z_k})|G] &= \boldsymbol{\Sigma_{u|k}} = \boldsymbol{\Sigma_{uu}} - \boldsymbol{\Sigma_{uk}}\boldsymbol{\Sigma_{kk}^{-1}}\boldsymbol{\Sigma_{ku}} \\ &= \boldsymbol{\Sigma_{uu}} - \boldsymbol{\Sigma_{uk}}\boldsymbol{\Sigma_{kk}^{-1}}\boldsymbol{\Sigma_{uk}^{T}} \end{aligned} \tag{2}$$

We estimate unknown statistics $\mathbf{Z_u}|G = \boldsymbol{\Sigma_{uk}}\boldsymbol{\Sigma_{kk}^{-1}} Z_k|G$, which can be interpreted as the unknown statistics associated with G being a weighted linear combination of known statistics of variants associated with G, with the weights reflecting the LD relationships among variants. We adopted the metric $r2pred = 1 - \boldsymbol{\Sigma_{u|k}}$ to measure the xQTL imputation accuracy, which represents the variances of unknown xQTL statistics $\mathbf{Z_u}|G$ explained by observed statistics $\mathbf{Z_k}|G$.

#### 2 IMPLEMENTATION FEATURES

XQTLImp is designed specifically to impute QTL statistics in association with thousands or even millions of molecular traits of different kinds of molecular types, such as gene expression (eQTL), DNA methylation (mQTL) and histone acetylation (haQTL), which differs from GWAS imputation, where only one phenotype is considered. The main challenge with taking a vast number of molecular traits into account is the computational cost in terms of time and memory due to the large number of associations that need to be considered during the imputation process. For example, the summary statistics of the GTEx cis-eQTL study [5] of brain amygdala contain 170 million eQTL associations genome-wide, with 23.6 thousand unique genes and 10 million unique variants. Thus, during implementation, we adopt several strategies to accelerate the imputation process. First, based on the observation that multiple traits, such as genes, might be located in the same LD, xQTLImp preserves a dynamic local LD matrix in memory to avoid redundant LD calculations while keeping the total memory cost in a low level. Second, xQTLImp allows the user to execute the imputation process in parallel using multiple threads with the *–num_threads N* parameter, during which associations on each chromosome will be broken into N chunks. Third, xQTLImp also allows users to specify genome regions to exclude using *–exclude or –exclude_file*, such as the complex HLA region, and to control the MAF lower bound of variants in the reference panel to achieve better efficiency. Fourth, users can also specify the chromosome number for chromosome-wide imputation using the *–chr* parameter, which is practically useful for taking advantage of high-performance clusters (by submitting chromosome-wide imputation jobs to different nodes). Fifth, users can adjust the window size around the molecular trait by using the *–window-size* parameter, to balance the performance accordingly. In addition, xQTLImp is also memory efficient by adaptively reading xQTL records and genotypes in the reference panel into memory and saving only the local LD matrix. The performance of xQTLImp on real datasets is shown in Table S2.

### 3 EQTL ANALYSIS OF ROSMAP DATASETS

The Religious Orders Study (ROS) and Memory and Aging Project (MAP) are two longitudinal cohort studies of aging that include deidentified clinical, neuropathological, brain RNA-seq and SNP array data for AD cases and controls [6]. In the ROSMAP study, mRNA was extracted from the homogenate of the dorsolateral prefrontal cortex and sequenced on the Illumina HiSeq platform. We downloaded the read count summary table, array-based genotype data and clinical tables from the Synapse platform (`https://www.synapse.org/#!Synapse:syn3219045`) with approval. The summary of gene expression levels covered 640 individuals and nearly 5,6000 genes. We used genotype data measured by Affymetrix GeneChip 6.0 at the Broad Institute, which covered 1,708 individuals and 636,236 SNPs.

#### 3.1 Genotype data process

We applied PLINK2 (v1.9beta) [7] and in-house scripts to perform rigorous subject and SNP quality control (QC) for the genotype dataset, which included the following: (1) remove subjects with call rate < 95%, (2) remove subjects with gender misidentification, (3) remove SNPs with genotype call rate < 95% (4), remove SNPs with Hardy-Weinberg equilibrium testing P value < 1E-6, (5) remove SNPs with test-mishap P value < 1E-9, (6) remove SNPs with minor allele frequency (MAF) < 0.05, (7) remove inbreeding subjects based on heterozygosity F score, (8) IBS/IBD filtering, and (9) remove population outliers using smartPCA [8]. After QC, 572,307 variants and 1,664 individuals remained for downstream analysis.
The genotype imputation process was performed on Michigan Imputation Server (`https://imputationserver.sph.umich.edu/index.html#!`), where Minimac3 was used for phasing and imputation [9], and 1000G phase3 was selected as the reference panel. Prior to imputation, we used the imputation preparation checking tool (`http://www.well.ox.ac.uk/~wrayner/tools/`) to perform external QC to fit the requirements of the imputation server. Variants with MAF $\geqslant 0.05$ and imputation quality score $R^2 \geqslant 0.3$ were used for downstream eQTL analysis.

#### 3.2 Gene expression data process

We applied stringent QC procedures for the ROSMAP RNA-seq dataset. The read count table was first transformed into a TPM (transcripts per million) table. Then, we used relative log expression (RLE) analysis, Spearman-correlation-based hierarchical clustering, and D-statistics analysis to remove sample outliers with problematic gene expression profiles [10]. Sample mix-ups were excluded by comparing the reported sex with the sex determined by the expression of the female-specific XIST gene and male-specific Y-chromosome gene RPS4Y1. TPM values were log transformed after adding a pseudocount, and quantile normalization was applied after excluding genes with low expression levels. SVA [11] was applied to adjust covariates of batch, age, gender, RIN, PMI and recognition levels (cogdx score).

##### 3.3 Cis-eQTL association analysis

After QC, 26,662 genes and 565,765 variants prior to genotype imputation and 6,330,993 post imputation of 292 individuals with both genotypes and RNA-seq data were used for cis-eQTL association analyses. MatrixEQTL [12] with the additive linear model was used for eQTL association analyses. SNPs within 1 Mb with the TSS of a gene were included.

#### 4 DATASET AVAILABILITY

To reproduce the experiments in this study, we list the datasets and links used for the analyses as follows:

- eQTL summary statistics of GTEx study [5]: GTEx portal (https://gtexportal.org/home/index.html).
- eQTL, mQTL, haQTL summary statistics of brain xQTL study [13]: Brain xQTL server (http://mostafavilab.stat.ubc.ca/xqtl/).
- Genotype and RNA-seq data of ROSMAP study [6] (in control use): Synapse platform (https://www.synapse.org/#!Synapse:syn3219045).
- Single cell eQTL summary statistics [14]: (https://genenetwork.nl/scrna-seq/)
- CD4+ T cell eQTL summary statistics for replication study [15]: (https://genenetwork.nl/cd4cd8eqtlbrowser/)

#### 5 ACKNOWLEDGMENT

The Genotype-Tissue Expression (GTEx) Project was supported by the Common Fund of the Office of the Director of the National Institutes of Health and by NCI, NHGRI, NHLBI, NIDA, NIMH, and NINDS. The summary statistics used for the analyses described in this manuscript were obtained from the GTEx Portal V7.

The results published here are in part based on data obtained from the AMP-AD Knowledge Portal (doi:10.7303/syn2580853). Study data were provided by the Rush Alzheimer's Disease Center, Rush University Medical Center, Chicago. Data collection was supported through funding by NIA grants P30AG10161, R01AG15819, R01AG17917, R01AG30146, R01AG36836, U01AG32984, and U01AG46152; the Illinois Department of Public Health; and the Translational Genomics Research Institute.

### 6 SUPPLEMENTARY FIGURES

##### Supplementary Figure S1

In this experiment, we first randomly masked certain percentage of summary statistics of eQTL from GTEx study v7 (brain Amygdala, $n = 88$) [5], haQTL and mQTL from the brain xQTL study (frontal cortex, $n = 433$ and 468, respectively) [13] on chromosome 1, with MAF $\geqslant 0.01$ for all variants. Second, xQTLImp was applied to impute the concealed statistics based on 1000G phase3 reference panel (EUR population, $n = 503$) [16].

For each type/column of QTL imputation, the first row (A1, B1 and C1) shows the correlation between the masked Z-statistics and imputed Z-statistics with the masked percentage set to 40% and r2pred cutoff set to 0.6. The color legend shows density of associations in a local area. The second row (A2, B2 and C2) shows the Pearson's correlations under various masking percentages and r2pred cutoffs. The third row (A3, B3 and C3) shows the recall rates, defined as the percentage of masked associations that can be recovered by imputation (with r2pred score larger than the cutoff). *–window_size* was set to 500kb, 500kb and 100kb for eQTL, haQTL and mQTL imputation respectively.

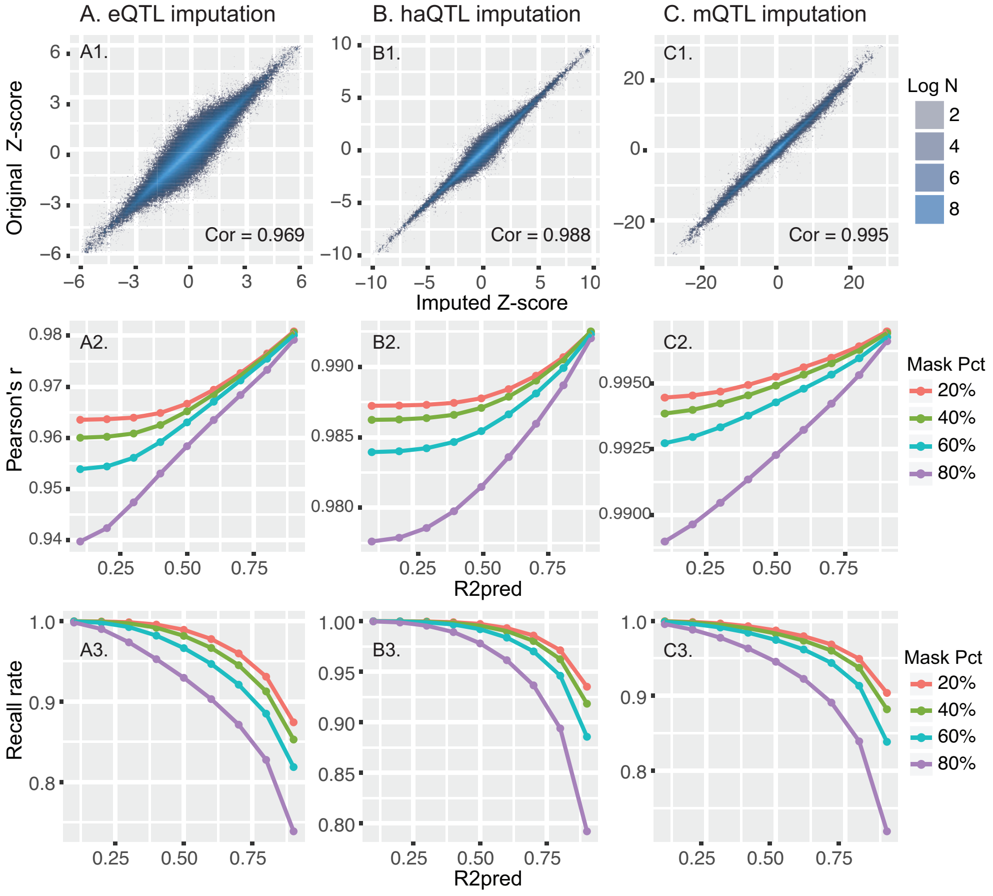

**Figure S1:** Accuracy of xQTLImp on eQTL, haQTL and mQTL statistics imputation.

##### Supplementary Figure S2

Multiple percentages (10% ~ 60%) of eQTL associations from GTEx [5] brain amygdala chromosome 1 ~ 22 were randomly masked prior to imputation, and Pearson's correlation coefficients between concealed Z-statistics and imputed Z-statistics on those hidden variant-gene pairs were calculated for measurement under different r2pred thresholds. The recall rate is defined as the percentage of masked associations with an imputation quality score (r2pred) larger than certain thresholds. Each boxplot represents the range of Pearson's r or recall rate among 22 chromosomes under the center r2pred threshold. The 1000G EUR population was selected as a reference panel with MAF $\geqslant 0.01$ during imputation, and the HLA region (chr6:25M-35M) on chromosome 6 was excluded.

To measure the effect of the random masking process, we repeated the random masking process on chromosome 1 five times, and the results of Pearson's r and recall rate show very small deviations (SDs range from $10^{-5}$ to $10^{-4}$, figure not shown).

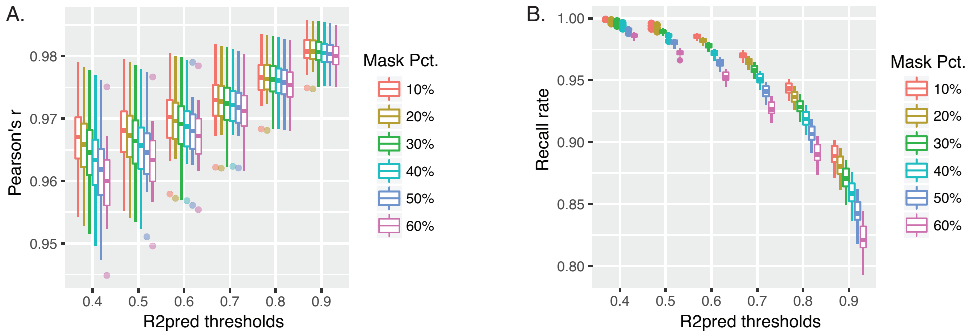

**Figure S2:** Imputation accuracy on different chromosomes.

##### Supplementary Figure S3

To test the imputation accuracy on lower MAF levels, two eQTL datasets, whose genotypes were called from WGS, brain amygdala ($N = 88$) and muscle skeletal ($N = 491$), from GTEx study v7 [5] were used for assessment. In the brain amygdala dataset, eQTL associations with variants having an $MAF > 0.1$ were used to impute associations with $0.05 \leqslant MAF \leqslant 0.1$. Pearson's correlation and recall rate were calculated in the same way as above, and the results are shown in Figure S3 panel A. The color in A2/B2 represented the density of associations in a small local area. To test the performance on an $MAF > 0.01$, we used the muscle skeletal dataset with a larger sample size. EQTL associations with variants having an $MAF > 0.05$ were used to impute associations with $0.01 \leqslant MAF \leqslant 0.05$. The results are shown in Figure S3 panel B. As shown, xQTLImp still achieved high accuracy with Pearson's r = 0.95 and 0.92, respectively, by setting a stringent r2pred cutoff as 0.9.

Notably, eQTL associations on chromosome 1 were used in this experiment, and the 1000G EUR population was selected as a reference panel with an $MAF \geqslant 0.01$ during imputation. The lower boundaries of MAF in the original GTEx study for brain amygdala and muscle skeletal were set to 0.056 and 0.01, respectively.

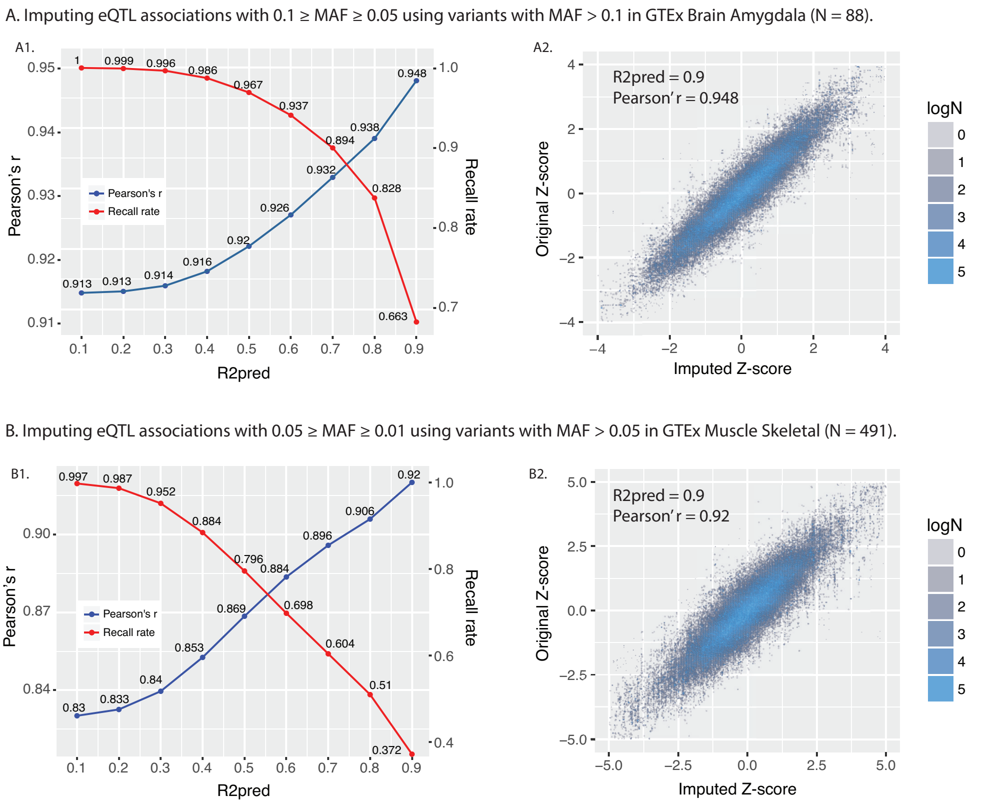

**Figure S3:** Imputation accuracy on lower MAF levels.

##### Supplementary Figure S4

To compare with the HMM-based genotype imputation framework, which is the gold standard when individual-level genotype data is available, we performed eQTL analyses in two ways. First, eQTL analysis was performed on genotyped variants only, and xQTLImp was adopted to impute the rest of associations against the 1000G phase3 reference panel. Second, individual-level genotypes were used for imputing the genotypes of the remaining variants on the 1000G phase3 reference panel by Minimac3 on Michigan Imputation Server [9], and then eQTL analysis was performed on those imputed variants (the traditional way). The Z scores of those imputed associations are shown in Figure S4, where the x-axis represents the Z scores obtained from the first way (xQTLImp imputation framework), and the y-axis represents the Z scores obtained from the second way (HMM-based imputation framework). Color represents the density of associations in a small local area. The result showed high consistency (Pearson's r = 0.996) between statistics from xQTLImp imputation and HMM-based imputation, indicating that the xQTLImp imputation framework based on summary statistics has similar accuracy as the HMM-based imputation framework based on individual-level genotype data.

Individual-level genotype data and RNA-seq data were obtained from the ROSMAP study [6] downloaded from the Synapse platform. Details of quality controls and eQTL mapping analysis can be found in section **"EQTL analysis of ROSMAP dataset"**. Chromosomes 1 ~ 22 were included for the analyses, excluding the HLA region (chr6:25M-35M) on chromosome 6, in this experiment. MAF lower bound was set to 0.05, and the other settings were their defaults.

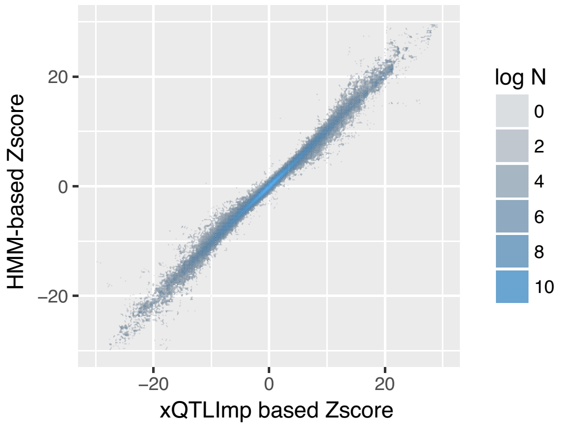

**Figure S4:** Imputation accuracy compared with HMM-based genotype imputation framework.

#### Supplementary Figure S5

An Upset plot was used to compare significant eQTL associations among six major cell types in PBMC [14], which was equivalent to a Venn diagram. The Upset diagram in panel A shows the original significant eQTL associations without eQTL imputation. The Upset diagram in panel B shows all significant eQTL associations after the eQTL imputation process. The left side bars show total number of significant eQTL associations found in each cell type, and the center bars show number of cell-type specific and common eQTL associations. We can see that a considerable amount of novel significant associations were rediscovered by our tool for all cell types. Upset plots were created with UpsetR [17].

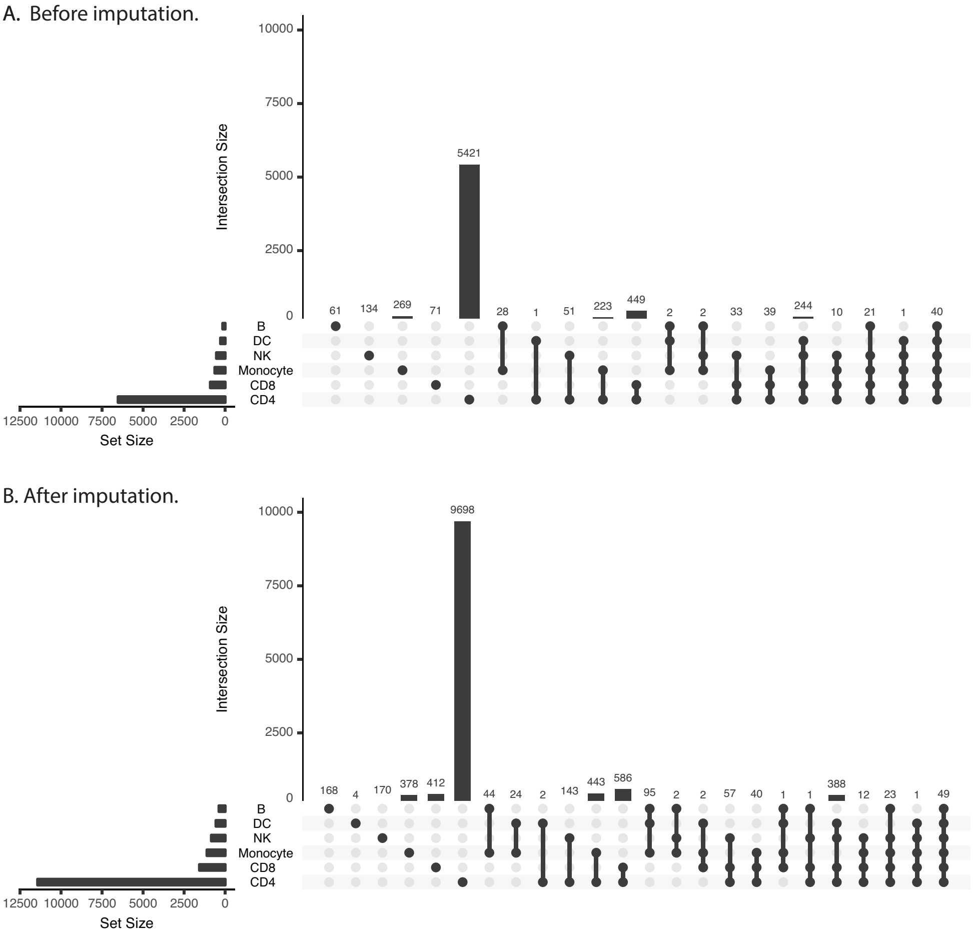

**Figure S5:** Upset diagram for significant eQTL associations in six cell types.

##### Supplementary Figure S6

Novel significant eQTL associations in CD4+ cell type (discovery study [14]) were replicated using an independent CD4+ T cell-type-specific eQTL dataset (N = 313, replication study [15]) . The lower bound of MAF of variants in the replication study was 0.05, and associations with an FDR < 0.5 were available to download. By limiting the comparisons to common genes and common variants between discovery and replication studies and setting FDR < 0.1 as a significance level in the replication study, we found that 85% of the novel significant associations in the discovery study can be replicated in the replication study, compared with 85.6% of the original significant associations that can be replicated. No significant difference (Fisher's exact test P value = 0.82) was found in the number of significant associations that can be replicated between original findings and our novel findings.

| | Replicated | Not Replicated | Replication percentage |
|---|---|---|---|
| Original significant eQTL-gene pairs | 2,483 | 419 | 85.6% |
| Novel significant eQTL-gene pairs | 369 | 65 | 85.0% |

Fisher's exact test
P value = 0.82

**Figure S6:** Diagram of two-way contingency table for Fisher's exact test.

### 7 SUPPLEMENTARY TABLES

**Supplementary Table S1**

Meanings of columns:

- Original sig. pairs: Number of significant eQTL-gene pairs in original eQTL summary dataset.
- Novel sig. pairs: Number of significant eQTL-gene pairs found by xQTLImp and not shown in original summary dataset.
- Original sig. eQTLs: Number of unique significant eQTLs (SNPs) in the original eQTL summary dataset.
- Novel sig. eQTLs: Number of unique significant eQTLS (SNPS) found by xQTLImp and not shown in original summary dataset.
- P cutoff (FDR $\leqslant$ 0.05): Significance P value cutoff corresponding with permutation-based FDR $\leqslant$ 0.05 in original study [14].

**Table S1:** Summary of novel significant eQTL associations found by xQTLImp compared with that in original single-cell eQTL datasets of six cell types.

| Cell type | Original sig. pairs | Novel sig. pairs | Original sig. eQTLs | Novel sig. eQTLs | P cutoff (FDR$\leqslant$ 0.05) |
|---|---|---|---|---|---|
| CD4 | 6489 | 4910 | 6320 | 4870 | 7.93E-06 |
| CD8 | 914 | 664 | 914 | 664 | 1.22E-06 |
| NK | 484 | 310 | 484 | 310 | 4.32E-07 |
| Monocyte | 383 | 479 | 372 | 473 | 6.97E-07 |
| DC | 155 | 229 | 155 | 229 | 5.08E-07 |
| B | 290 | 278 | 290 | 278 | 1.87E-07 |

##### Supplementary Table S2

In this experiment, approximately 5 million associations across chromosomes $1 \sim 22$ were used as input for each cell type, and 31.4 million associations on average were output for each cell type. The 1000G phase3 reference panel of the EUR population was used for LD calculation, with the following additional settings: MAF was set to 0.01, window size was set to 500Kb (250 Kb upstream and downstream from the center of a gene), and HLA region (chr6:25M-35M) was excluded.

**Table S2:** Running time and memory cost on single-cell eQTL datasets of six cell types.

| Cell type | Threads | Time(H:M:S) | Memory |
|---|---|---|---|
| CD4 | 20 | 3:55:29 | 3.58G |
| CD8 | 20 | 3:15:52 | 3.54G |
| NK | 20 | 3:57:20 | 3.56G |
| Monocyte | 20 | 3:46:25 | 3.59G |
| DC | 20 | 3:51:21 | 3.66G |
| B | 20 | 3:45:54 | 3.65G |

[4] Hanna Julienne, Huwenbo Shi, Bogdan Pasaniuc, and Hugues Aschard. Raiss: Robust and accurate imputation from summary statistics. *bioRxiv*, page 502880, 2018.
